# Lipid lowering effects of *Fucus vesiculosus* and *Phytolacca berry* extracts: An alternative approach of treating hypertriglyceridemia

**DOI:** 10.64898/2026.09.18.752646

**Authors:** Nazmul Hasan, Nazmul Ahsan, Tasnim Binte Ahmed, Anwarul Azim Akhand

## Abstract

Hypertriglyceridemia is commonly managed with lipid-lowering drugs such as statins; however, adverse effects, dose intolerance, and financial barriers may limit their use. Therefore, affordable and potentially safer plant-derived alternatives warrant investigation. This study evaluated the lipid-lowering effects of ethanol extracts of *Fucus vesiculosus* and *Phytolacca berry*, administered individually or in combination, in mice fed a butter-enriched diet for 12 weeks. Forty-eight mice were randomly assigned to six groups: normal control (Group 1), butter control (Group 2), statin control (Group 3), *Fucus vesiculosus* extract (Ext-1; Group 4), *Phytolacca berry* extract (Ext-2; Group 5), and the combined extracts (Group 6). Body weight, organ-to-body-weight ratios, serum lipid parameters, and histopathological changes were assessed. Group 3 showed the lowest final body weight, while Group 6 demonstrated a comparable body-weight trajectory. Organ-to-body-weight ratios varied among groups, with Group 3 generally showing the highest values and Group 6 comparatively lower ratios for several organs. Group 5 had the lowest total cholesterol concentration, followed by Group 3, whereas Group 6 showed cholesterol levels comparable to the normal control and statin groups. Notably, Group 6 exhibited the lowest serum triglyceride concentration (129.9 mg/dL). Histopathological examination revealed relatively mild hepatic alterations and limited vascular plaque formation in Group 6. The combined extracts demonstrated favorable effects on body-weight gain and serum lipid profiles, particularly triglycerides, with comparatively mild histopathological changes. Further mechanistic, dose-response, toxicity, and long-term studies are warranted before clinical application can be considered.

## Introduction

Lipids are hydrophobic organic molecules that are essential components of cell membranes, serve as energy stores, and participate in diverse cell-signaling pathways [1]. Major lipid classes include cholesterol and triglycerides (TGs). Cholesterol is an important biological molecule required for membrane structure and steroid hormone synthesis. Its biosynthesis begins with acetyl-CoA and acetoacetyl-CoA and proceeds through the formation of 3-hydroxy-3-methylglutaryl-CoA (HMG-CoA), which is subsequently reduced to mevalonate by HMG-CoA reductase [2]. Cholesterol is transported in the circulation primarily in association with lipoproteins and is synthesized mainly in the liver, whereas triglycerides are obtained from dietary sources and synthesized endogenously. TGs consist of glycerol esterified to three fatty acids and serve as an important form of energy storage [3]. A plasma TG concentration below 150 mg/dL is generally considered desirable, whereas 150–199 mg/dL is considered borderline high [4]. Elevated TG levels are associated with obesity, metabolic syndrome, type 2 diabetes, and increased cardiovascular risk [5, 6].

Fucus vesiculosus is a brown macroalga, commonly known as bladderwrack or seaweed, that contains a range of bioactive compounds, including phlorotannins, and has been reported to possess antioxidant, antimicrobial, and anti-inflammatory activities [7-12].

Phytolacca berry (Phytolacca spp.) has also been used in traditional medicine for various purposes, including anti-inflammatory and antiemetic applications [13,14]. Previous studies have suggested that extracts of Phytolacca species may influence metabolism and body-weight regulation [15-17].

Elevated blood cholesterol and triglyceride concentrations are important risk factors for cardiovascular and metabolic diseases, including fatty liver disease and type 2 diabetes [18]. Given the increasing burden of dyslipidemia in South Asia, including Bangladesh [19], additional affordable lipid-lowering interventions warrant investigation. Therefore, the present study evaluated the effects of F. vesiculosus and Phytolacca berry extracts, individually and in combination, on body weight, serum lipid profiles, organ weight ratios, and hepatic and cardiovascular histopathology in mice fed a butter-enriched diet.

## Material and Methods

### 2.1 Plant extract preparation

Fucus vesiculosus (Linnaeus, 1753; Aphia ID: 145548) extract (Ext-1) was prepared from fresh plant material, whereas Phytolacca berry (Linnaeus, Species Plantarum 441, 1753; IRMNG ID: 1071800) extract (Ext-2) was prepared from dried roots. After collection, the plant materials were thoroughly cleaned, dried in a cool, dry place protected from direct sunlight, and finely ground. The powdered samples were weighed and macerated in 65% ethanol (v/v) at a 1:10 (w/v) ratio for 3 weeks, with the containers shaken twice daily. To determine extract yield, 100 mL of each ethanol extract was evaporated, and the residual solid was collected and weighed. The mean yields were 9.69 ± 1.21 g/dL for F. vesiculosus and 9.72 ± 1.15 g/dL for Phytolacca berry.

### 2.2 The plant extract’s dose

The extract dose was selected in relation to the statin dose used in the experiment. A one-tenth dilution of each extract was used to limit potential excessive exposure while maintaining a proportional basis for comparison with the reference treatment. The rationale and exact administered dose should be reported explicitly in mg/kg body weight to facilitate reproducibility.

### 2.3 Animal maintenance

Forty-eight male Swiss albino mice, four weeks of age, were procured from the Department of Pharmacy, Jahangirnagar University, Bangladesh. The mice were randomly selected and housed in cages with wood-cob bedding (8 mice/cage). After one week of acclimation, the mice were divided into control and experimental groups.

### 2.4 Measurement of mice body weight

Each mouse was weighed every 2 weeks using an analytical balance, and body weight was recorded throughout the 12-week experimental period.

### 2.5 Measurement of the organs of mice

After 12 weeks, the mice were euthanized, and the abdomen was opened by a transabdominal incision. The heart, kidneys, liver, and spleen were carefully removed, cleared of adherent fat and connective tissue, and weighed.

### 2.6 Blood collections and assay of different serum parameters

After 12 weeks, the mice were euthanized, and blood was collected by cardiac puncture. Serum was separated by centrifugation at 4,000 rpm at 4 °C for 5 minutes and stored at −80 °C until analysis. Serum total cholesterol, triglycerides, HDL, and LDL concentrations were measured using a commercially available assay kit according to the manufacturer’s instructions (Human Diagnostic, Germany). Each serum sample was analyzed in duplicate, and the mean value was used for statistical analysis (Humalyzer 3000, Germany).

### 2.7 Slide preparation and histological analysis

The liver and heart were examined histologically. Sections were prepared using hematoxylin and eosin (H&E) staining [20] to assess tissue morphology and lipid-related changes, and Masson’s trichrome staining [21] to assess collagen deposition and fibrosis.

### 2.8 Statistical analysis

Statistical analyses were performed using SPSS version 24.0 (SPSS Inc., Chicago, IL, USA). Data are presented as mean ± SD. Differences among groups were assessed using one-way ANOVA followed by Bonferroni’s multiple-comparison test. A p value < 0.05 was considered statistically significant.

## Results

### 3.1 The combination of Ext-1 and Ext-2 limited body-weight gain

At baseline, the mean ± SD body weights of Groups 1–6 were 21.83 ± 2.98, 23.89 ± 3.88, 21.64 ± 3.14, 23.60 ± 2.43, 23.27 ± 2.07, and 21.99 ± 4.95 g, respectively. Body weight was measured at 0, 4, 8, and 12 weeks (Fig. 1A). After 12 weeks, the mean ± SD body weights were 52.91 ± 5.59, 59.98 ± 7.93, 46.74 ± 6.20, 53.94 ± 4.61, 57.53 ± 8.22, and 48.41 ± 8.38 g in Groups 1–6, respectively. Body weight increased progressively in all groups. Group 2 showed significantly greater weight gain than Group 1 (p < 0.05). In addition, weight gain in Groups 3, 4, and 5 was significantly lower than in Group 2 at the reported 4-, 8-, and 11-week time points (p < 0.05). Notably, the final body weight of Group 6 was similar to that of Group 3, indicating that the combined extract treatment was associated with a body-weight trajectory comparable to that observed with statin treatment (Fig. 1).

**Figure 1:**
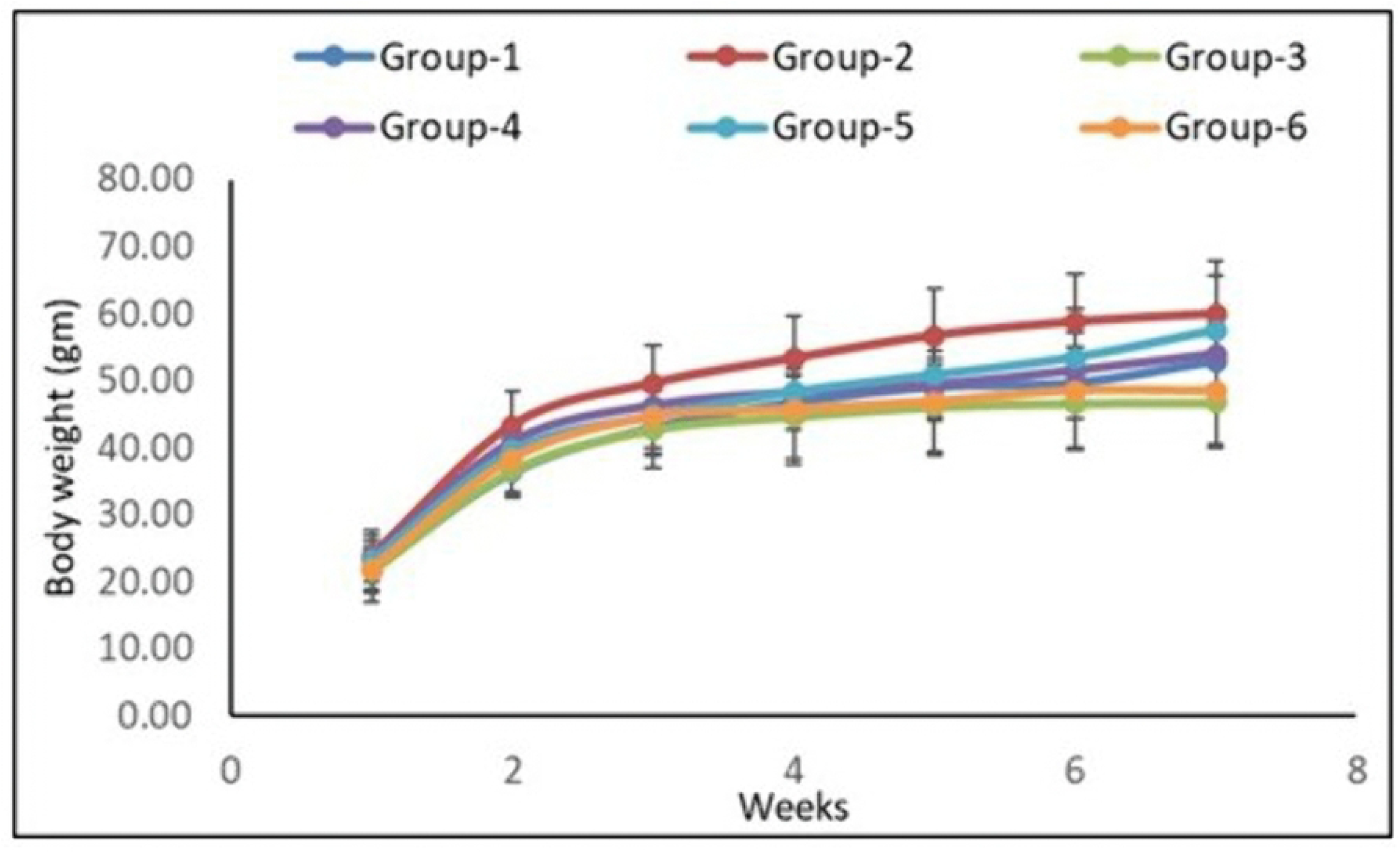
Changes in body weight over the 12-week experimental period. Group 2 showed the greatest weight gain, whereas Group 3 had the lowest final body weight; Group 6 showed a trajectory similar to Group 3.

#### 3.1.1 Effects of Ext-1 and Ext-2 on heart, liver, spleen, and kidney weight ratios

After 12 weeks, the weights of the liver, heart, kidney, and spleen were measured, and each organ weight was expressed relative to total body weight. The liver-to-body-weight ratio was highest in Group 3, followed by Groups 2 and 1, and was lowest in Groups 4–6, with Group 6 showing a significantly lower ratio than the other groups. The heart-to-body-weight ratio was highest in Group 3, followed by Groups 6, 4, 2, 5, and 1. The kidney-to-body-weight ratio was highest in Group 3, followed by Groups 1 and 6; Groups 4 and 5 showed similar ratios. The spleen-to-body-weight ratio was highest in Group 3, followed by Groups 2 and 1, whereas Group 6 had a higher ratio than Groups 4 and 5 but lower ratios than Groups 1–3 (Fig. 2).

**Figure 2:**
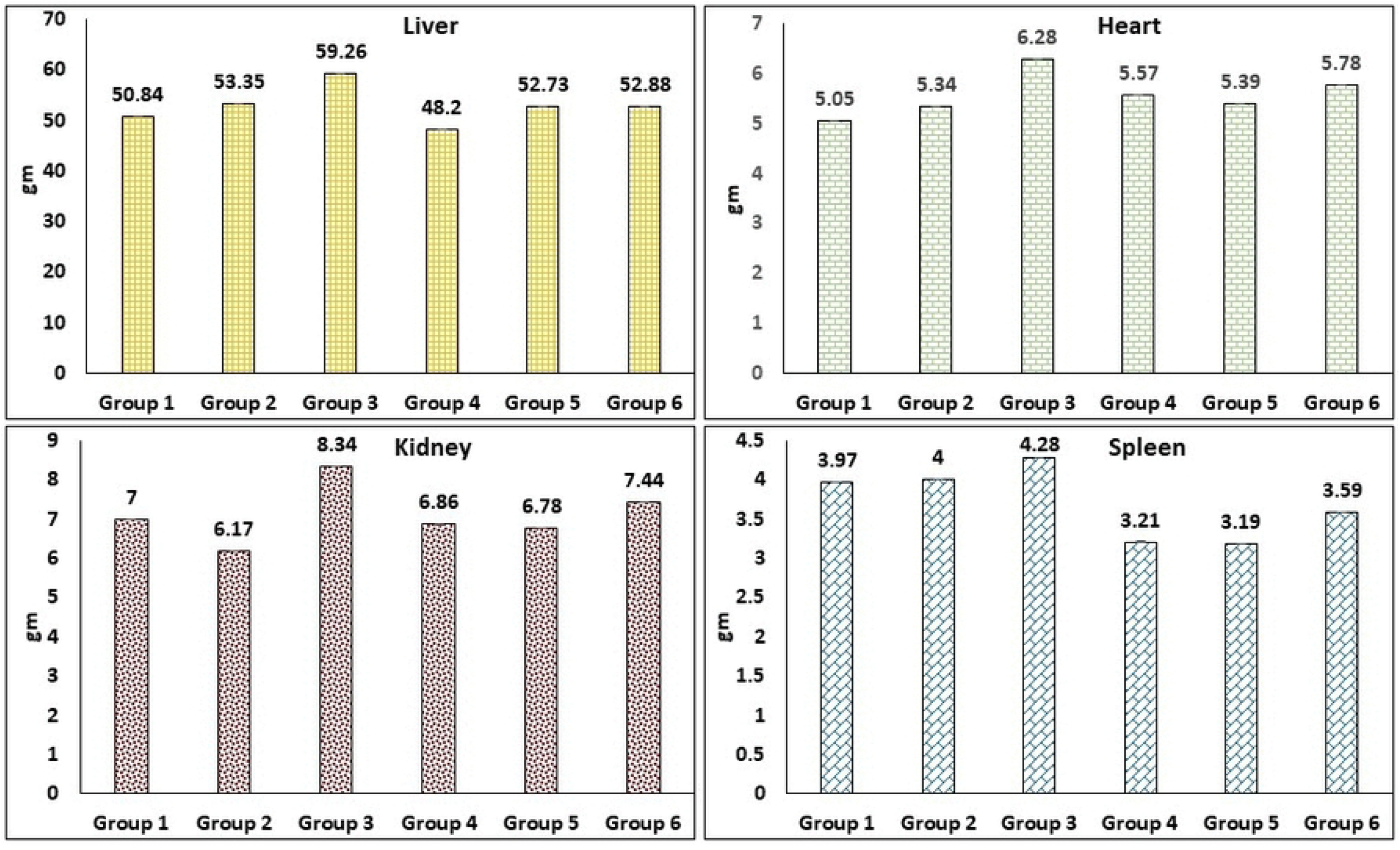
Organ weight-to-body-weight ratios after 12 weeks of treatment. The relative weights of the liver, heart, kidney, and spleen varied among the experimental groups.

### 3.2 Effects of Ext-1 and Ext-2 on serum total cholesterol

Serum total cholesterol concentrations were 211.4, 225.7, 211.0, 223.1, 208.4, and 213.4 mg/dL in Groups 1–6, respectively (Fig. 3A). The butter-fed control group (Group 2) had a higher cholesterol concentration than the normal control (Group 1), whereas Group 5 had the lowest value. Group 6 showed a cholesterol concentration close to those of Groups 1 and 3.

**Figure 3:**
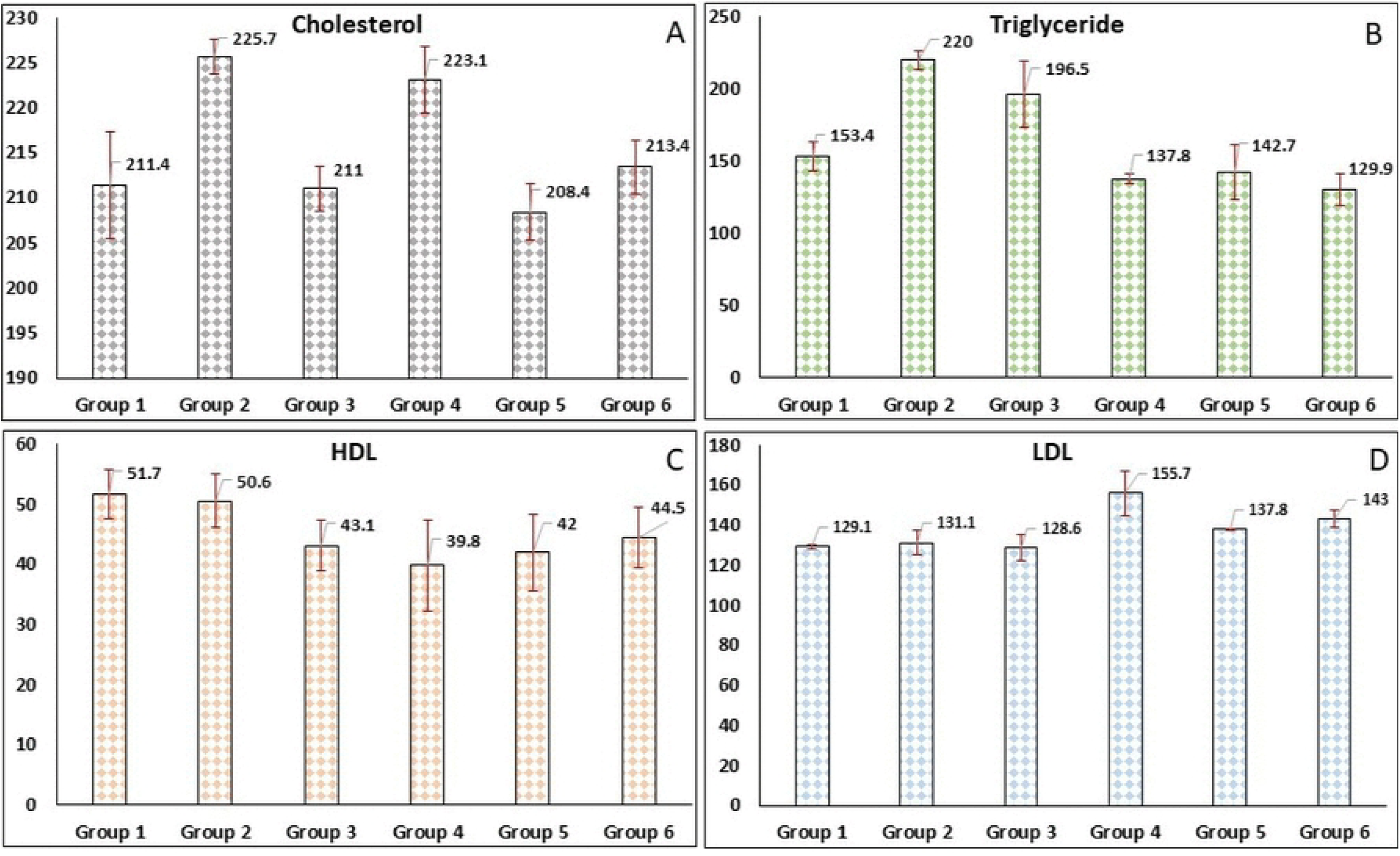
Effects of Ext-1, Ext-2, and their combination on serum lipid profiles. Group 2 showed higher total cholesterol and triglyceride concentrations than Group 1. Group 6 had the lowest triglyceride concentration and a total cholesterol concentration close to those of Groups 1 and 3.

#### 3.2.1 The combination of Ext-1 and Ext-2 reduced serum triglycerides

Serum triglyceride concentrations were 153.4, 220.0, 196.5, 137.8, 142.7, and 129.9 mg/dL in Groups 1–6, respectively (Fig. 3B). Groups 2 and 3 had higher concentrations than Groups 4–6. Group 6 had the lowest triglyceride concentration (129.9 mg/dL), which was below the commonly used 150 mg/dL threshold.

#### 3.2.2 Effects of Ext-1 and Ext-2 on HDL and LDL concentrations

HDL concentrations in Groups 1–6 were 51.7, 50.6, 43.1, 39.8, 42.0, and 44.5 mg/dL, respectively, whereas LDL concentrations were 129.1, 131.1, 128.6, 155.7, 137.8, and 143.0 mg/dL, respectively (Fig. 3C,D). Group 6 showed a higher HDL concentration than Groups 3–5 but remained below the level observed in Group 1. LDL was highest in Group 4 and lower in Group 5 and Group 6.

### 3.3 Combined Ext-1 and Ext-2 treatment was associated with relatively mild hepatic histopathological changes

Histopathological examination showed group-specific differences in hepatic steatosis, inflammation, ballooning, and fibrosis. Group 1 showed mild-to-moderate steatosis, mild inflammation, ballooning, and fibrotic changes. Group 2 showed mild-to-moderate steatosis, mild inflammation, and moderate ballooning. Group 3 showed mild steatosis and inflammation, with inflammation more pronounced than in the other groups, together with mild-to-moderate ballooning and mild fibrosis. Group 4 showed mild-to-moderate steatosis, mild inflammation, mild-to-moderate ballooning, and mild fibrosis. Group 5 showed mild steatosis, mild inflammation, moderate ballooning, and mild fibrosis. Group 6 showed mild steatosis, mild inflammation, mild-to-moderate ballooning, and mild fibrosis. Clustered steatosis was observed in Groups 1, 3, and 5, whereas diffuse steatosis was observed in Group 2. Groups 4 and 6 showed comparatively less severe overall hepatic changes (Figs. 4 and 5).

**Figure 4:**
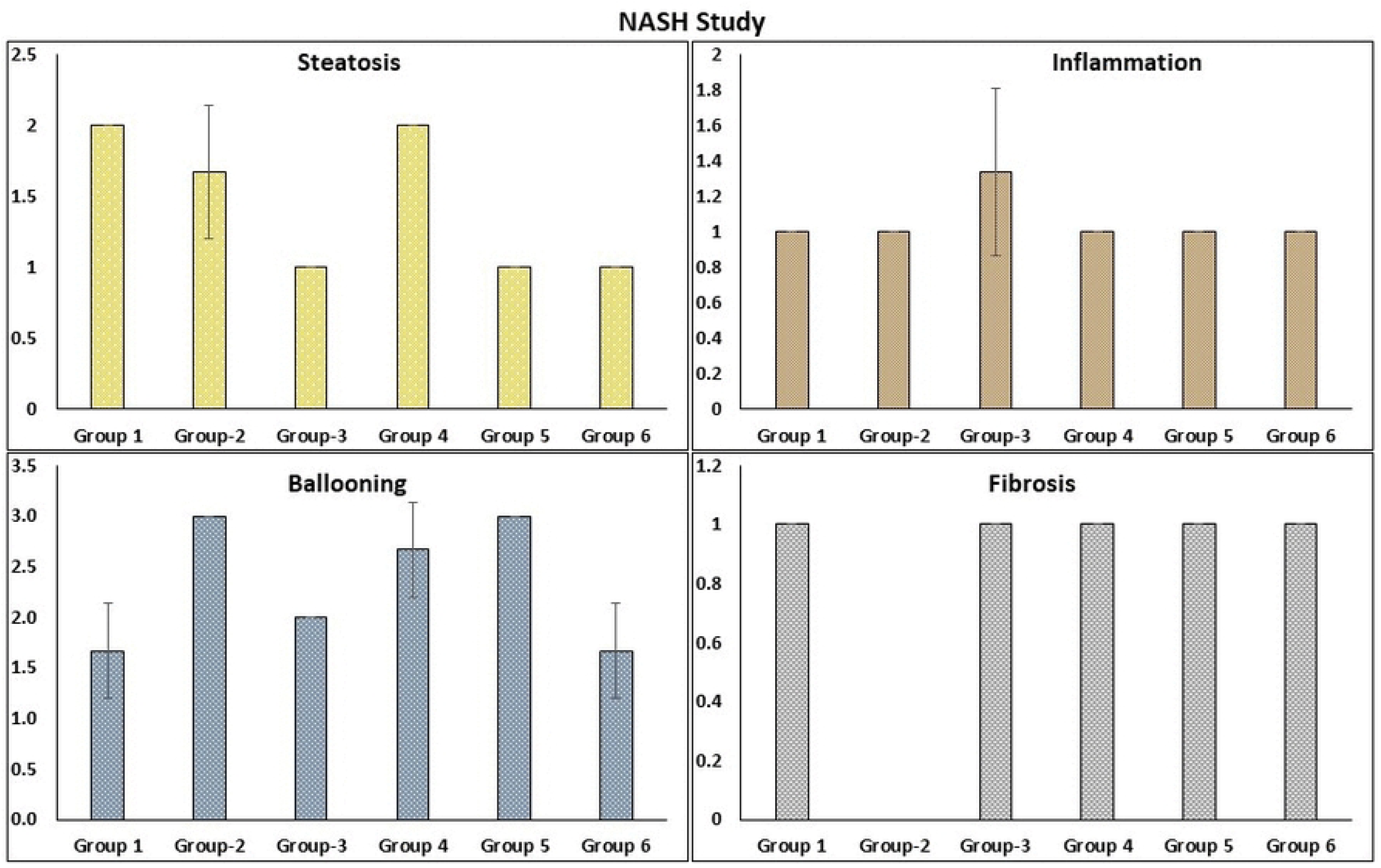
Representative liver histology after 12 weeks showing group-specific variation in steatosis, inflammation, ballooning, and fibrosis. Group 6 showed relatively mild changes across the assessed histological features.

**Figure 5:**
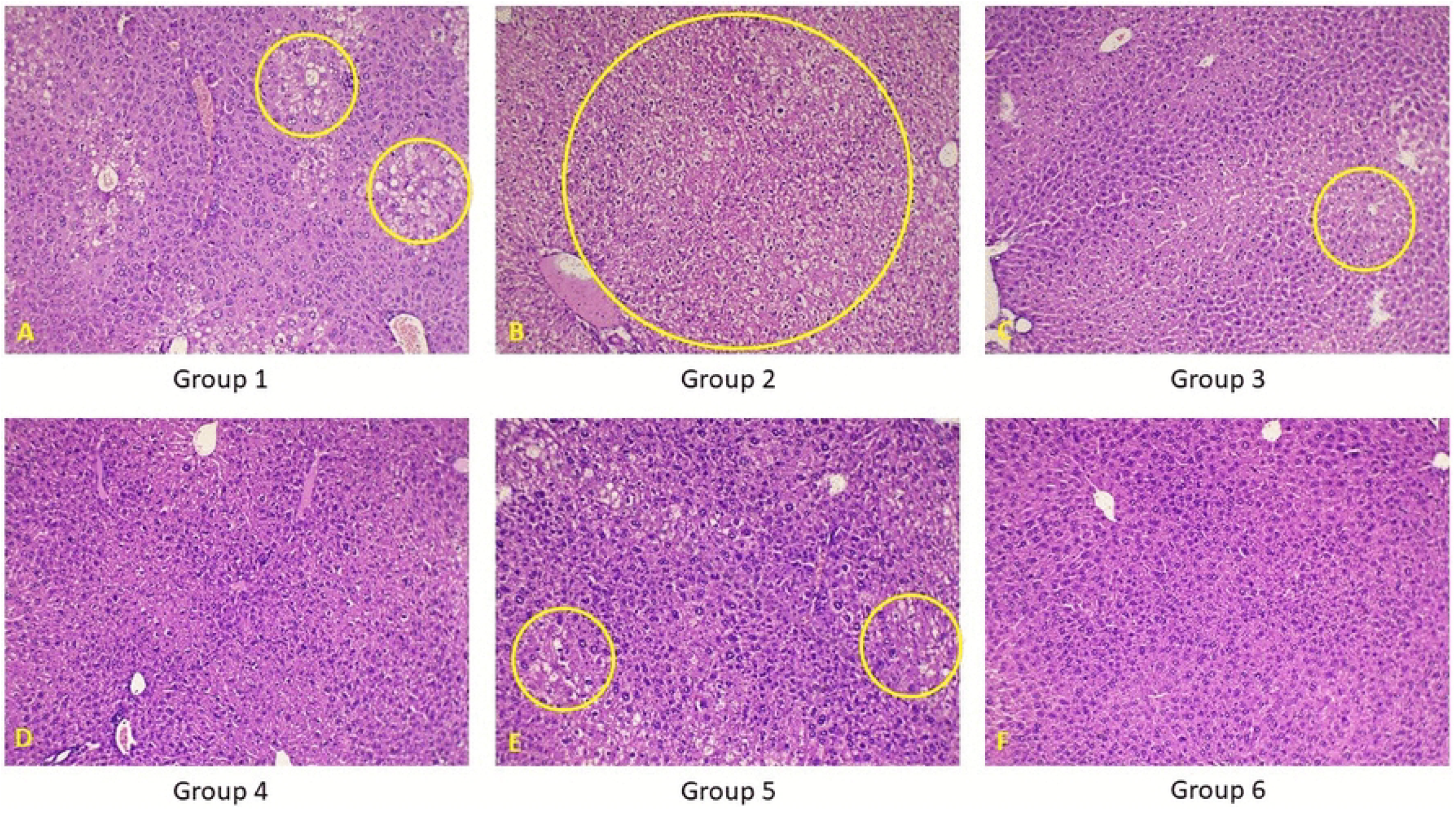
Comparative liver histology after 12 weeks. Clustered steatosis was observed in Groups 1, 3, and 5, whereas diffuse steatosis was observed in Group 2. Groups 4 and 6 showed comparatively less severe changes.

### 3.4 Combined Ext-1 and Ext-2 treatment was associated with less vascular plaque formation

Heart sections were examined for intimal thickening, plaque formation, calcification, fibrosis, and inflammation using H&E staining. Group 1 showed mild intimal thickening, plaque development, and inflammation. Group 2 showed mild intimal thickening, mild-to-moderate plaque development, and mild inflammation. Group 3 showed mild intimal thickening, plaque development, and inflammation. Groups 4 and 5 showed mild intimal thickening without evident plaque formation, inflammation, calcification, or fibrosis. Group 6 showed mild intimal thickening and the least plaque development among the groups, although mild fibrosis was observed in one mouse. Intimal thickening was present in all groups, whereas plaque formation was observed in Groups 1–3 and, to a lesser extent, in Group 6 (Figs. 6 and 7).

**Figure 6:**
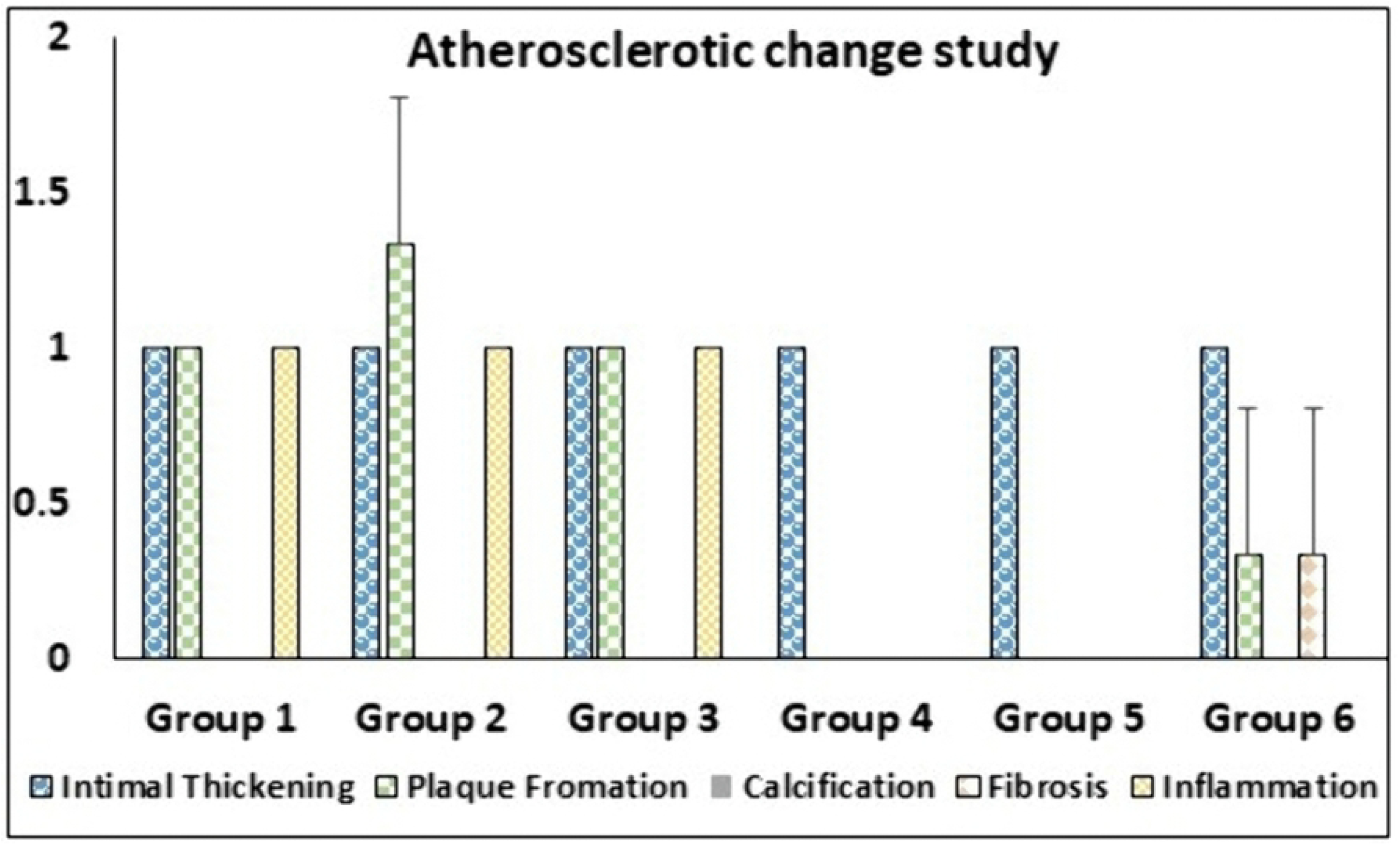
Representative heart histology after 12 weeks. Mild intimal thickening was observed in all groups. Plaque formation was mild in Groups 1 and 3, mild-to-moderate in Group 2, and minimal in Group 6. Inflammation was observed in Groups 1–3; no inflammation, calcification, or fibrosis was observed in Groups 4 and 5, whereas mild fibrosis was observed in Group 6.

**Figure 7:**
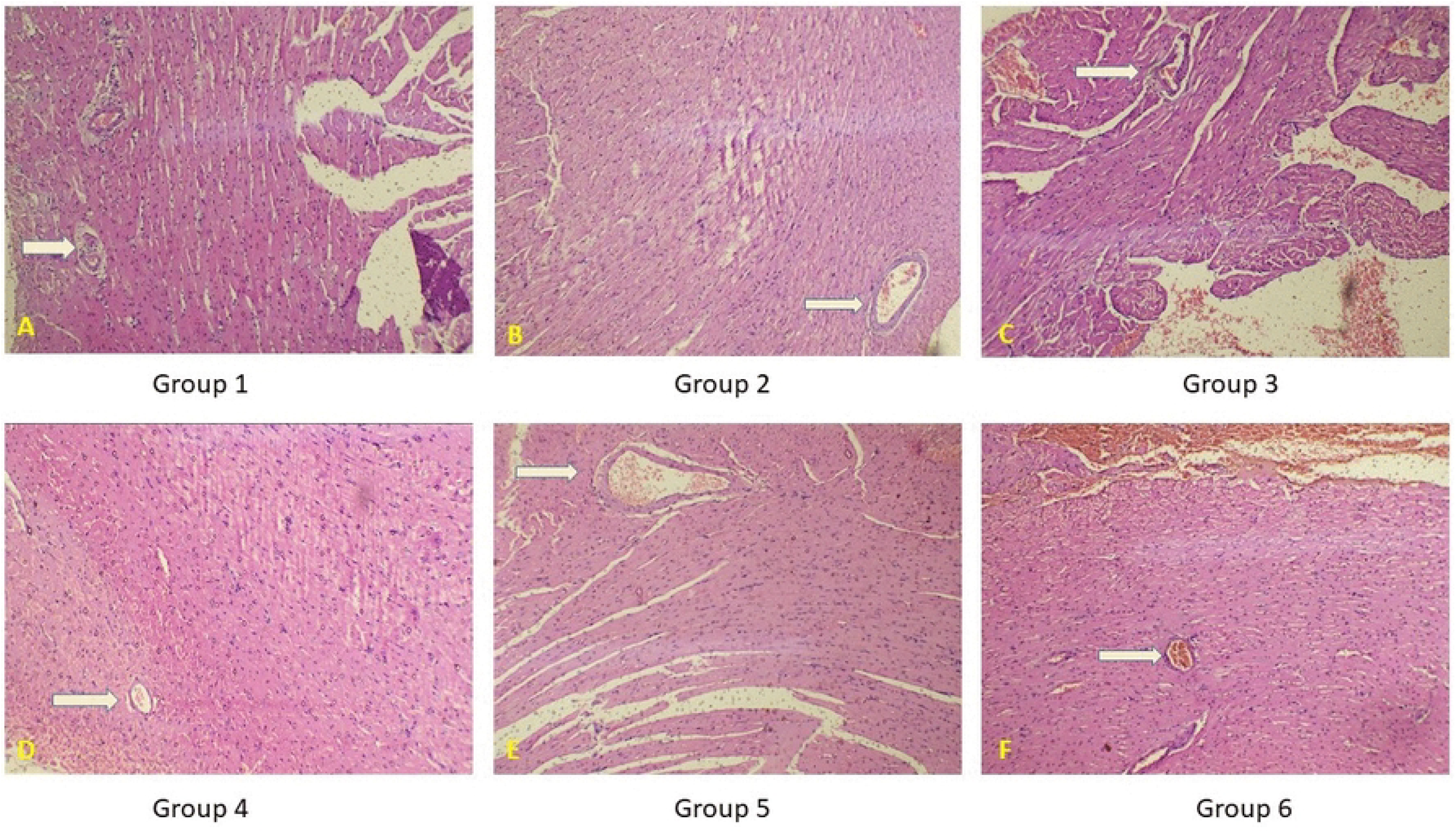
Comparative heart histology after 12 weeks. Mild intimal thickening was observed in all groups. Plaque formation was observed in Groups 1 and 3, was mild-to-moderate in Group 2, and was minimal in Group 6. Inflammation occurred in Groups 1–3, while mild fibrosis was observed in one mouse in Group 6.

Overall, Group 6 showed the most favorable triglyceride profile, with a serum triglyceride concentration of 129.9 mg/dL. Its total cholesterol concentration (213.4 mg/dL) was close to those of the normal control (211.4 mg/dL) and statin group (211.0 mg/dL). Histopathological examination also showed relatively mild hepatic changes and limited vascular plaque formation in Group 6. These findings identify the combined extract treatment as the most promising intervention among the experimental groups, although the available data do not establish mechanism, long-term safety, or clinical efficacy.

## Discussion

This study evaluated the effects of F. vesiculosus (Ext-1) and Phytolacca berry (Ext-2), administered individually or in combination, on body weight, serum lipid profiles, organ weight-to-body-weight ratios, and hepatic and cardiovascular histopathology in mice fed a butter-enriched diet. The most consistent finding was that the combined treatment (Group 6) produced the lowest serum triglyceride concentration and a body-weight profile similar to that of the statin group, while also showing comparatively mild histopathological changes.

Body weight increased progressively in all groups over the 12-week period, but the magnitude of gain differed among treatments. Group 2, which received the butter-enriched diet without lipid-lowering treatment, showed the greatest weight gain, consistent with the expected obesogenic effect of a high-fat diet [22, 23]. Groups 3–5 showed significantly lower weight gain than Group 2 at the reported time points (p < 0.05). Importantly, Group 6 had a final body weight close to that of Group 3. Thus, the combined extract treatment was associated with attenuation of diet-associated weight gain, although the present data do not demonstrate that the extracts directly regulate energy balance or adiposity.

The organ weight-to-body-weight ratios showed marked variation among groups. Group 3 generally had the highest relative organ weights, whereas Group 6 had a comparatively low liver ratio and intermediate heart, kidney, and spleen ratios. These observations suggest that the combined treatment did not produce the pronounced relative organ-weight increases observed in the statin group. However, organ-weight ratios alone are not sufficient to infer improved organ function or reduced hypertrophy; such conclusions would require biochemical, molecular, or functional assessments.

The serum lipid profile provides the strongest evidence for a beneficial effect of the combined extract. Total cholesterol was highest in Group 2 (225.7 mg/dL) and lowest in Group 5 (208.4 mg/dL), while Group 6 showed a value of 213.4 mg/dL, close to the normal-control and statin groups. More notably, Group 6 had the lowest triglyceride concentration (129.9 mg/dL), lower than the values observed in Groups 1–5. This result suggests that the combined extract may have a particularly favorable effect on triglyceride metabolism. Nevertheless, because statistical significance was not reported for all pairwise lipid comparisons, the magnitude of these differences should be interpreted descriptively rather than as evidence of superiority over statin treatment.

HDL and LDL responses were less favorable and less consistent than the triglyceride response. Group 6 had an HDL concentration of 44.5 mg/dL, which was higher than those of Groups 3–5 but lower than that of Group 1. LDL was highest in Group 4 (155.7 mg/dL), whereas Group 6 showed an intermediate value (143.0 mg/dL). These findings indicate that the lipid-lowering effect of the combined treatment was not uniform across all lipid fractions. Therefore, the principal apparent benefit of the combination in this study was reduction of triglycerides, with a comparatively modest effect on total cholesterol and no clear LDL-lowering advantage.

The histopathological findings provide supportive, but largely qualitative, evidence of tissue effects. Group 6 showed mild steatosis, mild inflammation, mild-to-moderate ballooning, and mild fibrosis, and its overall hepatic changes appeared less severe than those observed in several other groups. These findings are compatible with a potentially protective effect against diet-associated hepatic injury. However, the study does not provide quantitative histological scores or statistical comparisons for these features; therefore, claims of significant hepatoprotection should be avoided.

Similarly, vascular histology showed mild intimal thickening in all groups. Plaque formation was most pronounced in Group 2 and was absent in Groups 4 and 5, while Group 6 showed only minimal plaque formation. Mild fibrosis was observed in one mouse in Group 6. These observations suggest that the combined extract was not associated with extensive vascular pathology under the experimental conditions. Because the histological assessment was descriptive and plaque-related endpoints were not quantitatively analyzed, a cardiovascular protective effect cannot yet be established.

Taken together, the results indicate that the combined extract produced a favorable overall pattern, particularly with respect to triglyceride reduction and attenuation of body-weight gain. The combination should not, however, be described as definitively equivalent or superior to statin treatment because several endpoints lacked reported statistical comparisons, and the study was conducted in mice. The results are better interpreted as evidence that the combination warrants further investigation as a potential lipid-modulating intervention.

The greater apparent effect of the combined treatment than of either extract alone may indicate an additive or synergistic interaction between Ext-1 and Ext-2. Several mechanisms could plausibly contribute, including antioxidant and anti-inflammatory activity, altered lipid absorption, enhanced fatty-acid oxidation, modulation of adipogenic pathways, or effects on the gut microbiota. However, none of these mechanisms was directly examined in the present study. Future experiments should therefore test these hypotheses using molecular and biochemical endpoints, including relevant lipid-metabolism genes and proteins.

Several limitations should be acknowledged. The study used a single extract dose and a single treatment duration, and no dose-response relationship was established. Detailed biochemical markers of liver and kidney function, direct measures of adiposity, and comprehensive toxicity assessments were not reported. In addition, the histopathological findings were primarily descriptive, and mechanistic pathways were not investigated. Future studies should include multiple doses, longer follow-up, quantitative histopathological scoring, comprehensive toxicity assessment, and molecular analyses such as PPARγ, FAS, and SREBP-1c expression, together with investigations of gut-microbiota changes. Such studies will be important for determining reproducibility, mechanism of action, and translational potential.

## Conclusion

In conclusion, the combined F. vesiculosus and Phytolacca berry extracts were associated with reduced body-weight gain, the lowest serum triglyceride concentration among the experimental groups, and relatively mild hepatic and vascular histopathological changes in mice fed a butter-enriched diet. The triglyceride-lowering effect was particularly notable, while effects on total cholesterol, HDL, and LDL were more modest and variable. These findings suggest that the extract combination has potential as a plant-derived lipid-modulating intervention, but they do not establish clinical efficacy or safety. Dose-response, mechanistic, long-term toxicity, and translational studies are needed before the combination can be considered a therapeutic alternative to statins.

## Acknowledgments

This study is a part of the PhD research work of Nazmul Hasan who received partial financial support from UGC as PhD Fellowship

## Acknowledgments

This study is a part of the PhD research work of Nazmul Hasan.

## Ethical clearance

The study protocol was reviewed and approved by the Ethical Review Committee of the Faculty of Biological Sciences, University of Dhaka (Ref. No. 311/Biol. Scs.).

## Author Contributions

N.H.: Carry out the experiments, methodology, data curation, validation, analysis, visualization, and writing; N.A.: conceptualization, methodology, data validation, review and editing, and supervision; T.B.A.: validation of histopathology data and review; A.A.A.: conceptualization, methodology, data validation, review, editing, and supervision.

## Data Availability Statement

All data generated or analyzed during this study are included in this article. Further inquiries may be directed to the corresponding author.

## References

1. Fahy E, Cotter D, Sud M, Subramaniam S. Lipid classification, structures and tools. Biochimica et Biophysica Acta (BBA)-Molecular and Cell Biology of Lipids. 2011 Nov 1;1811(11):637–47.

2. Lin CJ, Lai CK, Kao MC, Wu LT, Lo UG, Lin LC, Chen YA, Lin H, Hsieh JT, Lai CH, Lin CD. Impact of cholesterol on disease progression. Biomedicine. 2015 Jun 1;5(2):7.

3. Jain BP, Pandey S, Goswami SK. Protocols in biochemistry and clinical biochemistry. Elsevier; 2024 Oct 16.

4. Pejic RN, Lee DT. Hypertriglyceridemia. The Journal of the American Board of Family Medicine. 2006 May 1;19(3):310–6.

5. Yuan G, Al-Shali KZ, Hegele RA. Hypertriglyceridemia: its etiology, effects and treatment. Cmaj. 2007 Apr 10;176(8):1113–20.

6. Choi W, Kang JH, Park JY, Hong AR, Yoon JH, Kim HK, Kang HC. Elevated triglyceride levels are associated with increased risk for major adverse cardiovascular events in statin-naïve rheumatoid arthritis patients: a nationwide cohort study. InSeminars in Arthritis and Rheumatism 2023 Dec 1 (Vol. 63, p. 152274). WB Saunders.

7. African Register of Marine Species [Internet]. Basel: Fucus vesiculosus Linnaeus, 1753; Odido M, Appeltans W, BelHassen M, Mussai P, Nsiangango SE, Vandepitte L, et al., editors [cited 2025 Dec 17]. Available from: https://www.marinespecies.org/afremas/aphia.php?p=taxdetails&id=145548.

8. Pereira J, Ribeiro PA, Santos AM, Monteiro C, Seabra R, Lima FP. A comprehensive assessment of the intertidal biodiversity along the Portuguese coast in the early 2000s. Biodiversity Data Journal. 2021 Oct 8; 9:e72961.

9. Prew ZS, Reddy MM, Mehta A, Dyer DC, Smit AJ. The African seaforest: a review. Botanica Marina. 2024 Oct 28;67(5):425–42.

10. Catarino MD, Silva AM, Cardoso SM. Phycochemical constituents and biological activities of Fucus spp. Marine drugs. 2018 Jul 27;16(8):249.

11. Golshany H, Siddiquy M, Elbarbary A, Seddiek AS, Kamal A, Yu Q, Fan L. Exploring Fucus vesiculosus phlorotannins: Insights into chemistry, extraction, purification, identification and bioactivity. Food Bioscience. 2024 Oct 1;61:104769.

12. Hill AF. The correct names of certain economic plants. Botanical Museum Leaflets, Harvard University. 1939 Jun 19;7(6):89–111.

13. Rogers GK. The genera of Phytolaccaceae in the southeastern United States. Journal of the Arnold Arboretum. 1985 Jan 1;66(1):1–37.

14. Mamedov N, Mehdiyeva NP, Craker LE. Medicinal plants used in traditional medicine of the Caucasus and North America. Journal of medicinally active plants. 2015 Jan 1;4(3-4).

15. Hauffe D. Archaeobotanical Analyses of the Winterville Mounds Site (22ws500) and Other Southeastern Ceremonial Complexes.

16. Marinas IC, Oprea EL, Geana EI, Luntraru CM, Gird CE, Chifiriuc MC. Chemical composition, antimicrobial and antioxidant activity of Phytolacca americana L. fruits and leaves extracts. Farmacia. 2021 Sep 1;69(5):883–9.

17. Kiran GR, Raju AB. Antiobesity effect of Phytolacca berry in rats. Env Exp Biol. 2014 Nov 5;12:95-.

18. Ruqaiya Hasan RH, Aisha Javaid AJ, Khan KR, Safia Malik SM. Electrolytes changes induced by weight loss herbal drugs Phytolacca Americana and Phytolacca Berry in hypercholesterolemic rabbits.

19. Hasan R, Khan KR, Kiran S. ROLE OF PHYTOLACCA AMERICANA AND PHYTOLACCA BERRY IN LIPID PROFILE ALLERATION IN HYPERCHOLESTROLEMIA INDUCED RABBITS ORYCTOLAGUS CUNICULUS. CANADIAN JOURNAL OF PURE AND APPLIED SCIENCES. 2012 Feb:1797.

20. Coleman R, Hayek T, Keidar S, Aviram M. A mouse model for human atherosclerosis: long-term histopathological study of lesion development in the aortic arch of apolipoprotein E-deficient (E0) mice. Acta histochemica. 2006 Dec 6;108(6):415–24.

21. Fujii T, Fuchs BC, Yamada S, Lauwers GY, Kulu Y, Goodwin JM, Lanuti M, Tanabe KK. Mouse model of carbon tetrachloride induced liver fibrosis: Histopathological changes and expression of CD133 and epidermal growth factor. BMC gastroenterology. 2010 Jul 9;10(1):79.

22. Lin S, Thomas TC, Storlien LH, Huang XF. Development of high fat diet-induced obesity and leptin resistance in C57Bl/6J mice. International journal of obesity. 2000 May;24(5):639–46.

23. Guimaraes VH, Lelis DD, Oliveira LP, Borém LM, Guimaraes FA, Farias LC, de Paula AM, Guimarães AL, Santos SH. Comparative study of dietary fat: Lard and sugar as a better obesity and metabolic syndrome mice model. Archives of Physiology and Biochemistry. 2023 Mar 4;129(2):449–59.

